# Nystagmus in patients with congenital stationary night blindness (CSNB) originates from synchronously firing direction-selective retinal ganglion cells

**DOI:** 10.1101/555011

**Authors:** Beerend H.J. Winkelman, Marcus H. Howlett, Maj-Britt Hölzel, Coen Joling, Kathryn H. Fransen, Gobinda Pangeni, Sander Kamermans, Hiraki Sakuta, Masaharu Noda, Huibert J. Simonsz, Maureen A. McCall, Chris I. De Zeeuw, Maarten Kamermans

**Author notes:** Correspondence to: M. Kamermans,. These authors contributed equally (Beerend H.J. Winkelman, Marcus H. Howlett, Maj-Britt Hölzel).

## Abstract

Congenital nystagmus, involuntary oscillating small eye movements, is commonly thought to originate from aberrant interactions between brainstem nuclei and foveal cortical pathways. Here we investigated whether nystagmus associated with congenital stationary nightblindness (CSNB) can result from primary deficits in the retina. We found that CSNB patients as well as an animal model (*nob* mice), both of which lack functional nyctalopin protein (NYX, nyx) in ON bipolar cells (ON-BC) at their synapse with photoreceptors, showed oscillating eye movements at a frequency of 4-7Hz. *nob* ON direction selective ganglion cells (ON-DSGC), which detect global motion and project to the accessory optic system (AOS), oscillated with the same frequency as their eyes. In the dark, individual ganglion cells (GC) oscillated asynchronously, but their oscillations became synchronized by light stimulation. Likewise, both patient and *nob* mice oscillating eye movements were only present in the light. Retinal pharmacological manipulations that blocked *nob* ON-DSGC oscillations also eliminated their oscillating eye movements, and retinal pharmacological manipulations that reduced oscillation frequency of *nob* ON-DSGCs also reduced oscillation frequency of their eye movements. We conclude that, in *nob* mice, oscillations of retinal ON-DSGCs cause nystagmus with properties similar to those associated with CSNB in humans. These results show that the *nob* mouse is the first animal model for a form of congenital nystagmus paving the way for development of therapeutic strategies.

## Introduction

Congenital nystagmus (i.e. involuntary repetitive uncontrolled eye movements) (1, 2) forms a heterogeneous group of eye-movement disorders with eye movements ranging from sinusoidal-like oscillations (pendular nystagmus) to highly asymmetrical repetitive eye movements (jerk nystagmus). Many patients suffering from congenital nystagmus also have a low visual acuity, which limits the quality of their daily lives. It is commonly thought to originate from aberrant interactions between brainstem nuclei and foveal cortical pathways. Decades of research have not uncovered the underlying pathophysiological mechanism (s) of congenital nystagmus or its associated reduced vision.

Our recent clinical research indicated that some types of congenital nystagmus might have a retinal origin (3). We examined 11 infant boys aged 2 months through 2 years who presented with a tonic downgaze of both eyes, a chin-up head posture and a rapid horizontal congenital nystagmus. These infants also had congenital stationary night blindness (CSNB), reduced visual acuity and mutations in either nyctalopin (*NYX*) or the L-type voltage-gated Ca-channel subunit 1F (*CACNA1F*) genes (3). These proteins are highly specific to the photoreceptor to ON-BCs synapse. NYX is located post-synaptically on the ON-BC dendrites (4, 5), whereas CACNA1F is expressed pre-synaptically at the photoreceptor synaptic terminal (6–9). Mutations in either of these genes abolish ON-BC activity (8–11).

Here we studied eye movements of *nob* mice, which lack nyx and are a well-established model for CSNB (7, 8). Strikingly, *nob* mice have a pendular nystagmus with a frequency similar to that found in young CSNB patients. Further analysis of retinal processing and eye movements shows that the oscillating eye movements are caused by a mechanism within the retina that induces synchronized oscillations in ON direction selective retinal ganglion cells (ON-DSGC) that project to the accessory optic system (AOS). Our results show that this form of congenital nystagmus has a retinal origin.

## Results

We reanalyzed video material of a group of young children (3 months – 2 years) with a disconjugate small-amplitude horizontal pendular nystagmus in combination with tonic downgaze (3). These patients had mutations in genes coding for proteins expressed at the synapse between photoreceptors and ON-BCs (3). We measured the oscillation frequency of their horizontal pendular nystagmus as 6.25 ± 0.63 Hz (n=3) (Fig S1). Because one of patients carried a mutation in the nyctalopin (*NYX*) gene, we analyzed the eye movements of *nob* mice, which also lack functional nyctalopin protein, and are a well-established model of CSNB (7, 8). wt mice (Fig 1A, blue lines) followed a horizontally moving vertical sinusoidal grating (100% contrast, spatial frequency 0.1 cycles/deg, velocity: 10 deg/s) with smooth eye movements and kept their eyes fixated when the stimulus stopped (grey bars). In contrast, the smooth eye movements of *nob* littermates (red lines) initially followed the same horizontally moving vertical grating, but could not maintain that movement and, they could not fixate their eyes when the stimulus stopped moving. Regardless of stimulus motion, they had spontaneous, small-amplitude oscillating horizontal eye movements (Fig 1A & B). Power spectral density plots show that *nob* eye-movement oscillation frequency was ~5 Hz, both during the stationary and the moving phases of optokinetic stimulation, whereas wt eye movements had no such peak (Fig 1C, red & blue line, respectively). On average the oscillation frequency to stationary gratings of *nob* eye movements was 5.17 ± 0.18 Hz (n = 17) (Fig 1C, blue line).

**Figure 1.**
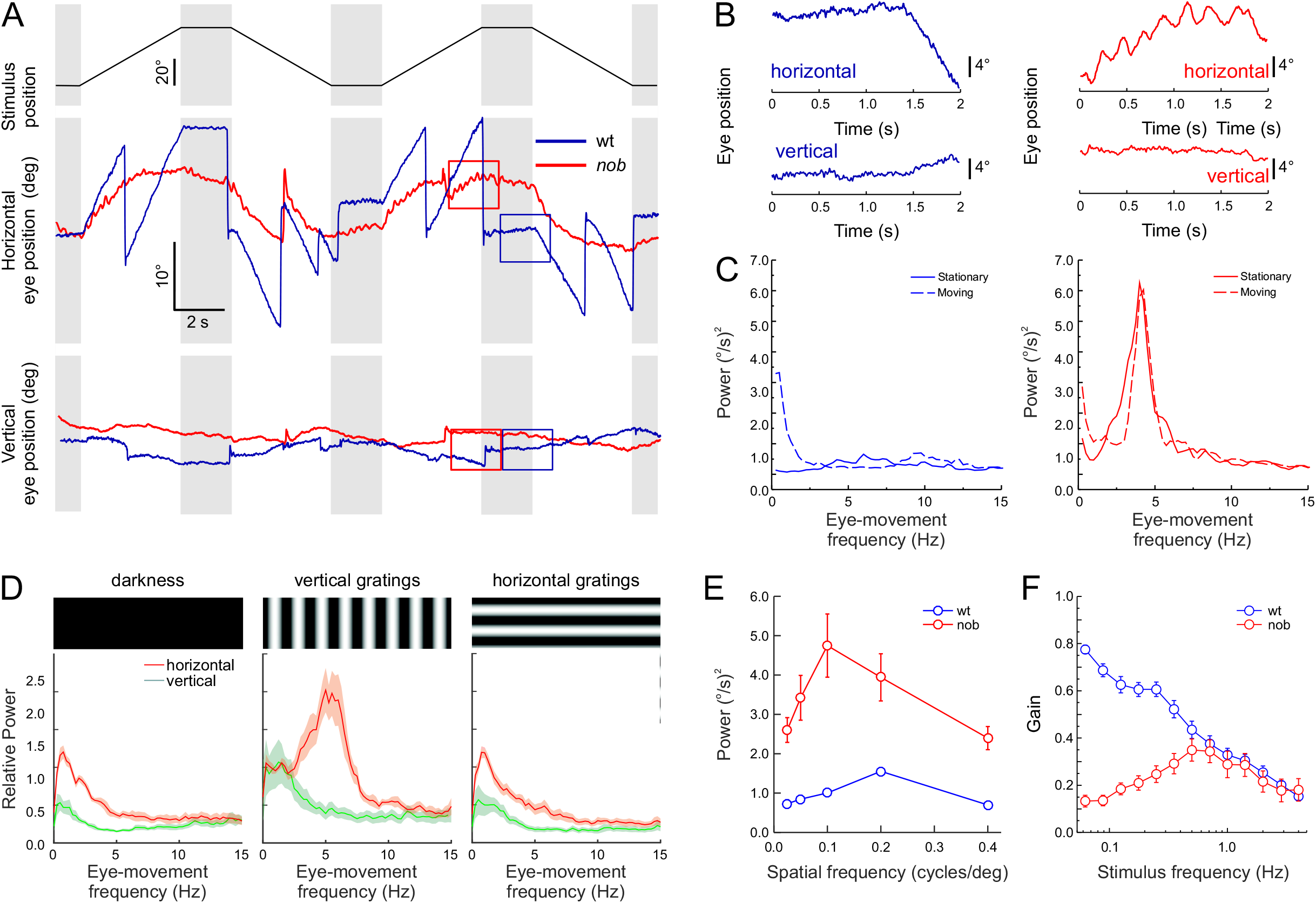
Horizontal eye movements are disturbed in *nob* mice. **A)** Schematic diagram of the timing of the stimulus used to evoke eye movements: 0.1 cycle/deg sine wave grating stimulus of 100% contrast moving at 10 deg/s (Top panel). Raw horizontal (middle panel) and vertical (bottom panel) eye movements are compared for wt (blue lines) and *nob* (red lines) mice. All *nob* mice tested behaved similarly (n = 9). **B)** Two second segments taken from the boxes in A on an expanded scale, show small amplitude oscillating eye movements in *nob* (right) and wt (left) mice. These 2 s segments taken from the traces of **A** (boxes) illustrate that wt mice made smooth eye movements in the presence of the moving contrast, whereas the eyes of *nob* mice oscillate in the horizontal direction. Both wt and *nob* mice vertical eye movements were generally absent. **C)** Power spectral density plots of the eye-movement velocity show a ~ 5 Hz oscillation frequency in *nob* mice (red) during moving and stationary stimuli, whereas the eyes of wt mice (blue) do not oscillate under those conditions. The peak in the power spectrum for the moving stimulus (0.5 Hz) in wt results from stimulus-induced eye movements. **D)** Relative power spectral density plots comparing *nob* eye movements (n = 9) in darkness (left panel), during vertical sine grating presentation (middle panel), and during horizontal sine grating presentation (right panel). A vertical grating with a spatial frequency of 0.1 cycles/deg effectively induced eye-movement oscillations while, a horizontal grating of the same spatial frequency did not. **E)** The relation between spatial frequency and power eye-movement oscillations in the same mice. A spatial frequency of 0.1 cycles/deg was the most effective. Oscillating eye movements are absent in wt mice (n = 9). **F)** Frequency-response plots of the OKR, tested by projecting a horizontal oscillating a dot pattern (peak velocity of 18.85 deg/sec) on a screen around the mouse. The eye-movement gain is expressed as the average amplitude of the eye–movement response divided by the amplitude of the stimulus. In wt mice (blue line; n = 11) the gain drops with increasing oscillation frequency. The eye-movement gain of n*ob* mice (red line; n = 9) deviate from wt at low frequencies.

By varying the spatial frequency and orientation of the stationary gratings, we determined that a vertical grating with a spatial frequency of 0.1 cycles/deg generated eye-movement oscillations with the strongest power (Fig 1D & E, red). Darkness and horizontally oriented gratings failed to induce significant eye-movement oscillations (Fig 1D). No vertical oscillating eye movements were detected (green).

Because the *nyx* mutation affects signaling by retinal ON-BCs (8–11), we asked whether *nob* mice show changes in their retinal output. We recorded activity from GC axons in the optic nerve (Fig 2A) and found that *nob* GCs showed oscillating burst-spiking behavior with a mean frequency of 4.79 ± 0.13 Hz (n = 46), which did not differ significantly from the mean frequency of their eye-movement oscillations (Students t-test, t = −1.53, df = 61, p = 0.13). These oscillations were absent in wt GCs (Fig 2A). Because of the wide spread oscillations across *nob* GCs, we hypothesized that retinal image motion signaling by DSGCs might be affected.

**Figure 2.**
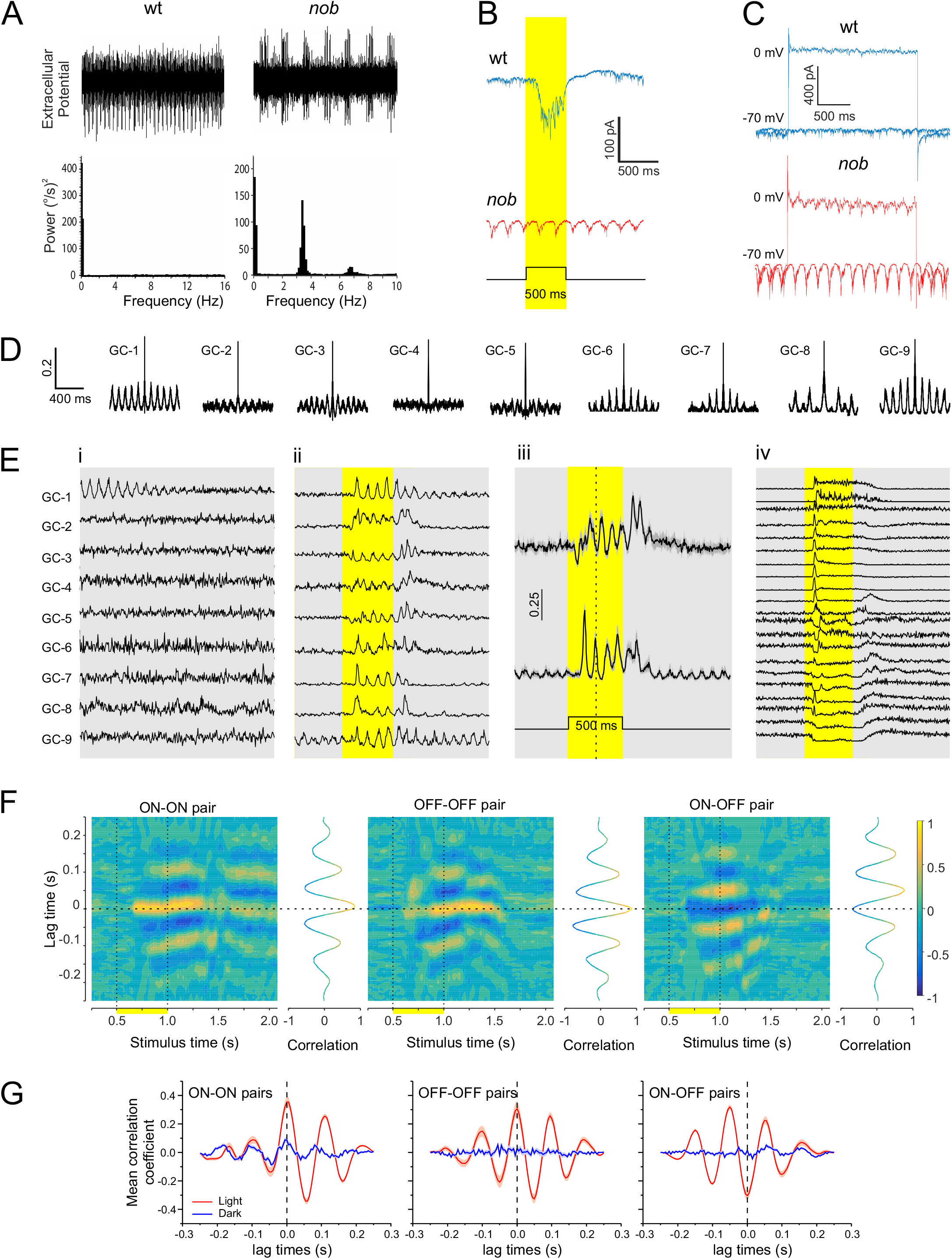
*nob* mice ganglion cell oscillations, asynchronous in the dark, are synchronized by light stimulation. **A)** Optic nerve recordings of spontaneous GC spiking activity in wt and *nob* mice after 30 mins of dark adaptation. The spontaneous activity of *nob* GCs shows oscillatory spiking patterns with a mean frequency of 4.79 ± 0.13 Hz (n = 46 isolated units). In this example, the GC fundamental frequency is 3.5 Hz. **B)** GFP-positive cells in wt/SPIG1^+^ mice show an increased inward current during a light flash, e.g., an ON response (blue trace). In *nob*/SPIG1^+^ mice, GFP-positive GCs lack a light evoked inward current and show oscillating inward currents. **C)** Inhibitory and excitatory currents in GFP positive wt/SPIG1^+^ (blue) and nob/SPIG1^+^ (red) GCs recorded under voltage clamp conditions (holding potential: 0 and −70 mV, respectively). In *nob* mice both excitatory and inhibitory inputs oscillate with a mean frequency of 4.87 ± 0.20 Hz (n = 38) and 4.25 ± 0.31 Hz (n = 8), respectively. **D-G)** Results based on GC spiking responses recorded on a MEA. **D)** Autocorrelations of 9 representative *nob* GCs from one retina in the dark. Periodic variations in their autocorrelations indicate spontaneous oscillatory activity. **Ei)** Each trace shows the mean normalized GC activity of 100 episodes of activity in the dark during the first 2 s of a 5 s window. All episodes were aligned to the first spike in the corresponding episode of GC-1 and then averaged. A clear oscillatory pattern is initially apparent for GC-1. In contrast, none of the other cells showed a similar pattern suggesting that each *nob* GC had a frequency and/or phase, independent of GC-1. **Eii)** Mean normalized activity of the same *nob* GCs in response to a 500 ms light flash (yellow shading). Note the presence of oscillation in all GCs, indicating that their phase was reset by the light flash. **Eiii)** Mean (± SEM) light evoked oscillations of GCs could be separated into two clusters that were in antiphase with each other. One cluster showed a decrease in spike rate just after light onset identifying them as OFF-GCs (top; n=10), whereas the other cluster responded with a delay and in antiphase suggesting they were ON-GCs (bottom; n = 13). **Eiv)** Mean responses of 21 wt GCs to the same stimulus used in Eii-iii show no oscillatory activity evoked by the light flash. **F)** Short-time cross-correlations between mean responses of representative *nob* ON-ON, OFF-OFF and ON-OFF GC pairs during the same light stimulation. Line graphs flanking the heat maps (3D plots) show the cross-correlation for a time window of 250 ms, advancing in 25 ms steps. After light onset, and around zero lag time, peak-positive correlation coefficients are found for *nob* ON-ON and OFF-OFF GC pairs and peak-negative correlation coefficients for the ON-OFF GC pair. **G)** Mean cross-correlations of all *nob* ON-ON (n = 78), OFF-OFF (n = 45) and ON-OFF (n = 130) GC pairs for *nob* GCs shown in **Eiii**. Two time windows are illustrated. The blue line indicates a window immediately prior to light onset (250 ms-500 ms), the red line a window during the light flash (680 ms – 930 ms). Panels **F** and **G** show that the oscillations are poorly synchronized in the dark before the light flash and that light induces the synchronization which slowly dissipates again in the dark after the light flash.

Image-motion signals are transmitted from the retina to the brain via two general pathways. ON-DSGCs, which are most sensitive to low temporal frequencies, low image velocities, and signal global motion and project to the AOS (12–15). In contrast, ON/OFF-DSGCs, which have a broad temporal frequency-response range, are mostly implicated in relative-motion detection and project mainly to the lateral geniculate nucleus (LGN) and superior colliculus (SC) (12–15). To determine which of these two pathways is affected most in *nob* mice, we recorded the frequency response curves of their eye movements induced by sinusoidally oscillating dot patterns with a constant peak velocity (Fig 1F). The frequency-response relation of *nob* (red symbols) and wt eye movements (blue symbols) differed, but only in the low frequency range, consistent with a malfunction in ON-DSGCs, suggesting these GCs are predominantly involved in generating the nystagmus phenotype.

To test this idea, we recorded excitatory and inhibitory currents in *nob* ON-DSGCs, using *nob* mice crossed and backcrossed with SPIG1^+^ reporter mice where ON-DSGCs coding for upward image motion are GFP labeled (16, 17). In contrast to wt ON-DSGCs, *nob* GFP^+^ ON-DSGCs were visually non-responsive and both their excitatory and inhibitory currents oscillated with a mean frequency of 4.87 ± 0.20 Hz (n = 38) and 4.25 ± 0.31 Hz (n = 8) respectively (Fig 2B & C). The oscillation frequencies of *nob* eye movements, GC burst-spiking, ON-DSGC excitatory and inhibitory currents did not differ significantly from each other (ANOVA: F = 1.018, df = 93, p = 0.40) indicating that a retinal oscillator, pre-synaptic to *nob* ON-DSGCs, drives these oscillations (Fig S2A).

Using a multi-electrode array (MEA), we recorded the activity of *nob* GCs extracellularly and assessed their light-evoked responses and oscillatory activity. *nob* ON-DGCs do not respond to light stimuli and this makes it impossible to identify these neurons in MEA recordings. In the dark, many *nob* GCs showed periodic variations in their auto-correlations (Fig 2D), indicating that they oscillated. We never observed such oscillations in wt GCs in either the dark or after a light flash, (Fig 2Eiv). To examine whether *nob* GCs oscillated synchronously, we selected nine oscillating *nob* GCs and in the dark aligned the activity of GC-2 through GC-9 such that their first spike in each of one-hundred 2 s periods, spaced 3sec apart, corresponded with the first spike fired by GC-1 during the same period. The aligned traces were averaged (Fig 2Ei) and the averaged traces show that the spike patterns of GC-2 through GC-9 are unrelated to GC-1 indicating that their rhythmic firing differed in frequency and phase from the oscillations of GC-1. We also observed that the oscillations in the mean response of GC-1 reduced over time, suggesting that the frequency and phase of the spontaneous GC oscillations varied slowly over time. In contrast, a brief light flash synchronized oscillatory activity across all nine GCs (Fig 2 Eii). In the light, their averaged traces of 100 repetitions (Fig 2Eii) showed clear oscillation patterns, indicating that light stimulation acted to phase-reset their activity and produced synchronized oscillations.

We found two clusters of *nob* GCs that oscillated in antiphase (Fig 2Eiii). In one cluster, *nob* GCs responded to the light flash with a reduction in firing rate, identifying them as OFF-GCs. In the other cluster, GCs mean firing rates remained unchanged during and after a light stimulation, suggesting they were ON-GCs (all *nob* are visually unresponsive). During the light flash, the oscillatory activity of *nob* GCs synchronized within each cluster, as revealed by their short-time cross-correlations (Fig 2F). Both ON-ON GC and OFF-OFF GC pairs showed multipeak correlograms with a positive coefficient around zero lag time after light onset, indicating synchronized oscillatory firing. An ON-OFF GC pair showed a similar multipeak correlogram, but now had a negative correlation coefficient around zero lag time, indicating that these GCs oscillated in antiphase. Fig 2G shows the mean cross-correlations for all ON-ON, OFF-OFF and ON-OFF GC pairs before (blue) and during (red) the light flash. This behavior was found in all four retinas tested.

To establish a causal relationship between eye movement and GC oscillations in *nob* mice, we pharmacologically blocked or modified the excitatory inputs to the GCs and measured both GC oscillations and eye-movement oscillations. Blocking excitatory AMPA and NMDA inputs *in vitro* with a cocktail of 50 μM CNQX (50 μM DNQX) and 10 μM D-AP5 eliminated all *nob* GC oscillations (Fig 3Ai – iii). Similarly, intraocular injections of a similar cocktail *in vivo*, eliminated GC oscillations in the optic nerve (data not shown), as well as eye-movement oscillations (Fig 3Aiv). As we found two groups of *nob* GCs that oscillated in antiphase, these oscillations presumably arise from a common presynaptic source, likely the AII amacrine cells (AC). The AII ACs drive both ON- and OFF-GCs with opposite sign and are known to generate intrinsic oscillations in mice with photoreceptor degeneration (rd1 mouse) (18). In rd1 mice, the oscillation frequency of AII ACs can be decreased by blocking glycine receptors with the antagonists strychnine (STR) (18). Similarly, in *nob* retina, application of STR reduced both ON-DSGC oscillation frequency (Control: 4.30 ± 0.36 Hz; STR: 2.60 ± 0.20; n = 5; Paired Students t-test, t = 5.01, df = 4, p = 0.007) (Fig 3Bi – iii) and the eye movement oscillation frequency (Control: 5.38 ± 0.125 Hz; STR: 2.99 ± 0.13; n = 4; Paired Students t-test, t = 6.48, df = 3, p = 0.007) (Fig 3Biii & iv). Finally, since the oscillations in AII ACs also depend on the activity of an M-type potassium current (18), application of 30 μM linopiridine hydrochloride (LP), a blocker of M-type K-current (19–21), led to a decrease in oscillation frequency of the ON-DSGCs from 5.71 ± 0.53 to 2.83 ± 0.40 Hz (n = 6; Paired Students t-test, t 7.45, df = 5, p = 0.0007) and a decrease eye-movement oscillation frequency from 5.00 ± 0.00 to 1.75 ± 0.50 Hz (n = 3; Paired Students t-test, t = 6.48, df = 2, p = 0.023) (Fig 3C). For extended statistics of these pharmacological experiments see Fig S2B.Together, these results indicate that: 1) the AII ACs are critically involved in generating the oscillations in *nob* GCs; 2) there is a causal relation between the synchronous *nob* ON-DSGC oscillations and their eye-movement oscillations and 3) the oscillator driving the oscillating eye movements is located in the retina.

**Figure 3.**
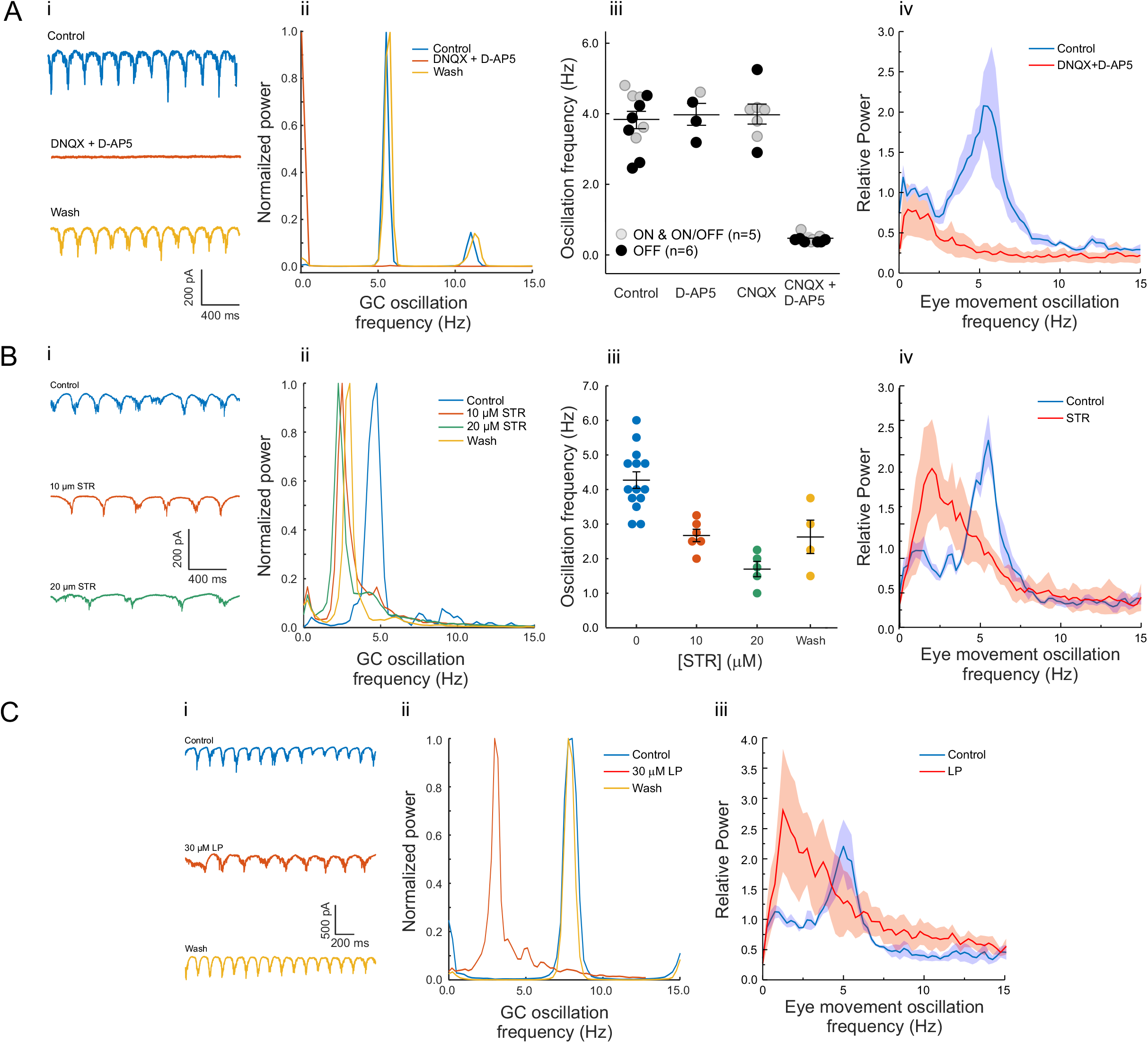
Pharmacological manipulations block of excitatory and inhibitory inputs in *nob* retina block or modify both GC and eye movement oscillations. **Ai)** Oscillating excitatory currents of *nob* ON-DSGCs (top) are blocked by a cocktail of50 μM DNQX and 10 μM D-AP5 (middle) and return after washout (bottom) (n=5). **Aii)** The power spectral density plot of the data in **Ai** shows that DNQX/D-AP5 eliminates the 5 Hz oscillating excitatory current. This was found in all *nob* GCs tested (n = 5). **Aiii)** CNQX or D-AP5 administered alone do not block excitatory current oscillations, whereas their combination blocks excitatory current oscillations in *nob* ON-, OFF- and ON/OFF-GCs. **Aiv)** Oscillating eye movements in *nob* mice (n = 5) are blocked by intravitreal injections of DNQX/D-AP5 (red; control: blue). **Bi & Bii)** Strychnine (STR) reduces the oscillation frequency of *nob* ON-DSGCs excitatory currents. **Biii)** Population data show that STR consistently reduced the oscillation frequency and that the oscillation frequency partly recovered after washout. **Biv)** Mean (± SEM) power spectral density plots (n=4) show that intraocular injection of STR lowered the oscillating frequency of the eye movements in *nob* mice. **Ci)** Bath application of LP shifted the oscillation frequency of *nob* ON-DSGCs excitatory currents to lower frequencies. **Cii)** The power spectral density plots of the ON-DSGC shown in **Ci**, show a shift in peak oscillation frequency to lower frequencies. This was found in all *nob* GCs tested (n = 6). **Ciii)** Mean (± SEM) power spectral density plots (n=3) show that intraocular injection of LP lowered the frequency of oscillating eye movements in *nob* mice. All GC recordings were done in the dark. For the experiments shown in Aiv, Biv and Ciii the stimulus was a stationary sinusoidal grating with a spatial frequency of 0.1 cycles/deg and 100% contrast.

## Discussion

This study reveals for the first time, a pathophysiological mechanism for a specific form of congenital nystagmus and shows that its origin is retinal. We propose the following mechanism (Fig 4 & S3). In wt mice, ON-DSGCs, which are sensitive to low-velocity global image motion (22), respond coherently when an image moves across the retina. Their coherent output reflects the direction and speed of global image movement: i.e. the retinal slip signal. This signal forms the input to the AOS where it induces compensatory eye movements that stabilize the image on the retina. In *nob* mice, the network behaves quite differently. *nob* ON-DSGCs are non-responsive to light stimuli, and thus cannot detect image motion, which may underlie the absence of a well-developed optokinetic response (OKR). In addition, *nob* ON-DSGCs oscillate spontaneously, which we show in this paper induces a pendular nystagmus.

**Figure 4.**
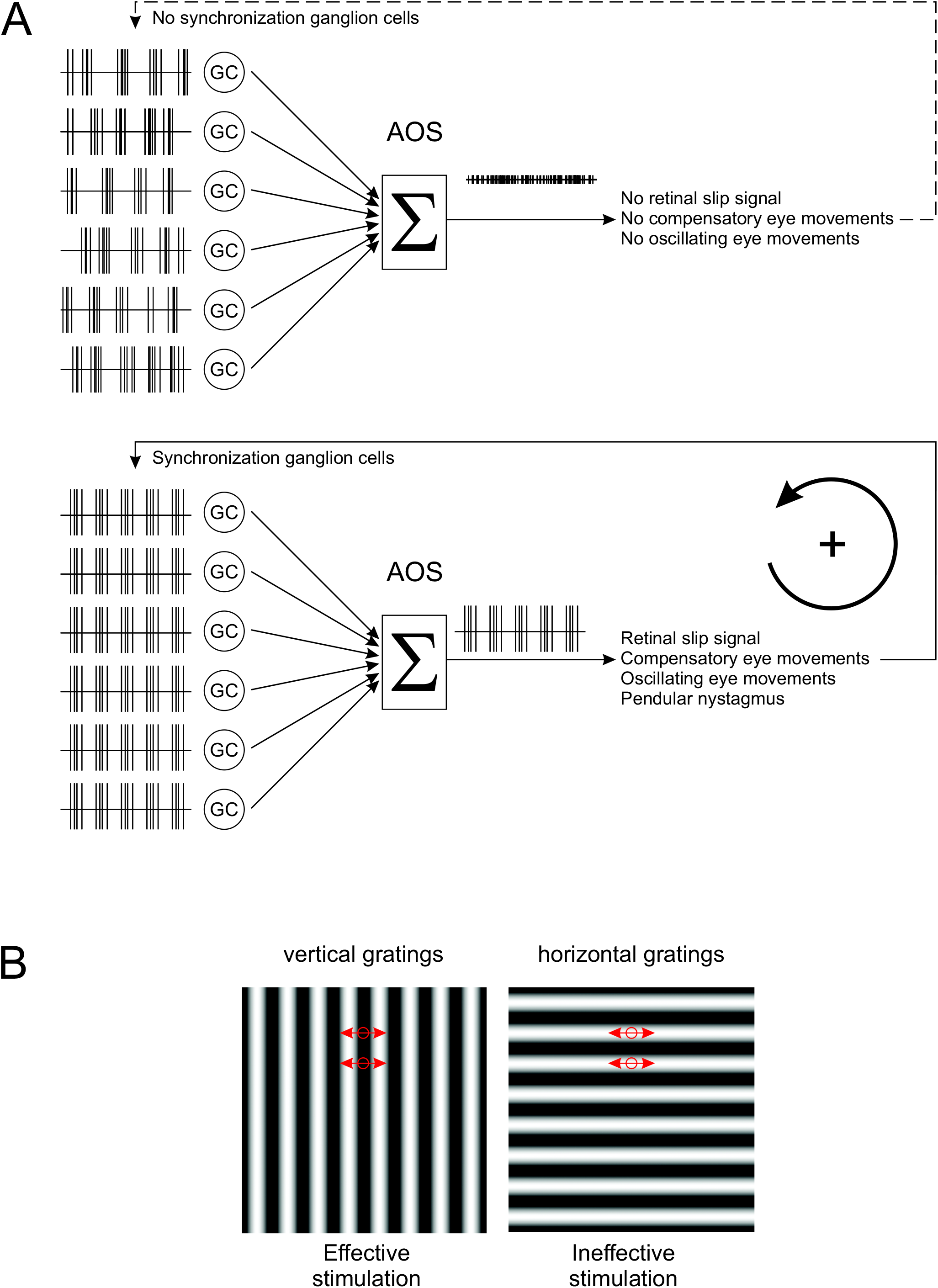
Model for the generation of night blindness-associated congenital nystagmus. **A)** wt ON-DSGCs (GC) signal global image motion to the AOS. The ON-DSGC signals are integrated (Σ) and a compensatory eye movement is induced. In the dark, *nob* ON-DSGC spiking activity oscillates but these oscillations are asynchronous. The integrated inputs to the AOS from asynchronous oscillating *nob* ON-DSGCs cannot generate a significant signal to trigger an eye movement. In the presence of a stimulus containing contrast, the oscillations of *nob* ON-DSGCs are synchronized and their integrated input to the AOS oscillates and generate an eye movement manifesting itself as a pendular nystagmus. **B)** A vertical grating oscillating horizontally over the retina will effectively activate GCs since the stimulus will change within the receptive field of the GC (Circle). A horizontal grating oscillating horizontally over the retina will be ineffective in inducing a response in retinal neurons since the stimulus will not change within the receptive field of the neurons (Circle).

We propose that the origin of these oscillations are the AII ACs for the following reasons. Firstly, the source of the oscillations is presynaptic to GCs. Second, AII ACs contain a membrane potential dependent oscillator, consisting of a fast sodium channel, and both a fast and a slow (M-type) potassium channel (18).When the AII AC membrane potential is outside its normal working range due to altered synaptic input, the AII ACs start to oscillate (18). Finally, the oscillations of ON- and OFF-GCs are driven in antiphase, which AII ACs will do since they drive ON- and OFF-GCs with opposite sign.

How could light stimulation synchronize the oscillations in the GCs? Since the oscillator in AII ACs is an intrinsic feature of the AII ACs, they could oscillate rather independent of each other leading to asynchronous GC oscillations as we find in the dark. We hypothesize that global light stimulation depolarizes the membrane potential of the AII ACs via cross-over inhibition driven by the OFF BCs. This will phase reset and synchronize the AII ACs and subsequently the GCs they project to. The consequence is that despite the absence of a classical visually evoked response in *nob* ON-DSGCs, their oscillations synchronize when the retina is stimulated with a global light stimulus. The combined activity of the synchronized oscillating *nob* ON-DSGCs is sent to the AOS where it is interpreted as an oscillating retinal slip signal. In response, oscillating compensatory eye movements are generated: i.e. pendular nystagmus emerges. On the other hand, when the oscillations of *nob* ON-DSGCs are asynchronous, such as in the dark, the combined activity will not oscillate and will not evoke pendular nystagmus.

Once initiated, oscillating eye movements over an image should induce retinal activity that phase resets the retinal oscillator (AII ACs), and maintains the synchronous oscillations of ON-DSGCs and pendular nystagmus (Fig 4). This represents a self-maintaining loop. Consistent with this idea, a horizontally oriented grating fails to induce oscillating eye movements, because horizontal eye movements over a horizontal grating appears as a stationary stimulus to GCs (Fig S2C). This will not modulate retinal activity and will not synchronize GC oscillations and pendular nystagmus is absent (Fig 1D).

The suggestion that AII ACs are the source of the oscillations implicates that all ON-DSGCs oscillate. If, this is the case, why is nystagmus in the CSNB patients and *nob* mice only horizontal when ON-DSGCs are tuned for image movement in three directions: naso-temporal, upward and downward (14). One of the simplest explanation is that the retinal slip signals generated by synchronously oscillating ON-DSGCs tuned for upward and downward motion cancel each other. In contrast, the naso-temporal signal is not canceled as there is no, or only a minor, opposing ON-DSGCs types (14).

In the present paper we discuss the origin of a specific form congenital nystagmus, i.e. spontaneous small amplitude involuntary oscillating eye movements. This pathological condition is distinct from the OKR in which the eyes follow the global movement of the stimulus followed by a fast reset saccade. The OKR, sometimes referred to as optokinetic nystagmus, is often reduced or absent in nystagmus patients. However, the conclusion that congenital nystagmus arises as a consequence of a mere absence of the OKR is unwarranted, since for example mice with mutations in FRMD7 (23) have no horizontal OKR but do not develop a pathological nystagmus.

It has been suggested that congenital nystagmus originates from a disruption in the interaction of the subcortical optokinetic pathways (AOS) and cortical foveal pursuit system (1). In contrast, our findings show that the cause of congenital nystagmus associated with CSNB lies within the retina in the afoveate mouse. Since the optokinetic systems of afoveate and foveate animals differ (24), generalizing our findings in *nob* mice to humans with CSNB requires caution. That said, given the balance of our results, we proposed that the *nob* mouse model is relevant to human CSNB patients for the following reasons. First, the nystagmus in young CSNB patients (Fig 1S) and *nob* mice are identical (Fig 1). Second, the photoreceptor to ON-bipolar cell synapse is highly conserved across mammals and even across vertebrates (25). Third, mutations in proteins specific for this synapse lead to the same CSNB phenotype in mice and humans (26), suggesting similar underlying retinal mechanisms. Forth, starburst amacrine cells (SAC), which are fundamental for generating direction selective responses of GCs (23), are found in both the mouse and primate retina (27). Finally retrograde tracing experiments reveal retinal projections to the AOS in both mice and primates (28). Moreover, it should be noted that even if the retinal projections to the AOS in human would not mediate physiological information about global image motion, they most likely carry an oscillation signal in CSNB patients that could induce oscillating eye movements, as are present in both the patients and the *nob* mice.

From a therapeutic point of view, our results suggest that the primary pathogenic condition in patients with this form of congenital nystagmus occurs in the retina in the form of synchronized oscillations of GCs. Therapeutic interventions aimed to de-synchronize the oscillating GCs in the retina might serve to eliminate nystagmus.

## Material & Methods

### Animals

All animal experiments were carried out under the responsibility of the ethical committee of the Royal Netherlands Academy of Arts and Sciences (KNAW) acting in accordance with the European Communities Council Directive of 22 July 2003, (2003/65/CE) or the University of Louisville Animal Care and Use Committee. Wild type mice, *Mus musculus*, were obtained from Janvier labs. *nob* mice were obtained from the McCall lab (University of Louisville, Louisville, KY, USA) and SPIG1^+^ mice were obtained from the Noda lab (National Institute for Basic Biology, Okazaki, Japan). Since the *nob* mutation is X-linked only male animals were used. For wt SPIG1^+^, both male and female animals were used. All mice were kept in a C75BL/JRj background. The age of the animals ranged from 4 weeks to 1 year.

### Eye movement recordings

#### Surgical preparation

Prior to the start of experiments, adult animals were equipped with a head-fixation pedestal, an aluminum bit with an integrated magnet, attached to the parietal bones of the skull using dental cement (Super Bond, Sun Medical). Surgery was performed under general isoflurane/O_2_ anesthesia and topical anesthesia (bupivacaine). Analgesia was offered by subcutaneous injection of metacam (2 mg/kg). A recovery time of at least 2 days subsequent to pedestal surgery. During experiments, the animal was placed head-fixed in the experimental setup using a custom-made adapter, which allowed panoramic vision.

#### Optokinetic stimulator

For the optokinetic experiments two experimental setups used. In the first system a modified Marquee 9500 CRT projector (100 fps) generated a large panoramic visual surround (545 nm) onto three large back-projection screens (1.47 × 1.18 m; Stewart Filmscreen) that were placed around the animal to create a combined field of view 270 × 77.5 deg. The average luminance of the visual stimulus was 10 cd/m^2^. Mickelson contrast of the grating stimuli was about 90%. In the second setup two Benq XL2420t high performance monitors (120 fps, gamma-1.745) were placed in V-formation around the animal and the closest distance between the screen surface and the mouse head was 16.5 cm. Screen dimensions were 56.9 × 33.8 cm (combined field of view: 240° × 50°). Mickelson contrast of the grayscale grating stimuli approached 100%. The average luminance of the sine grating stimuli was 71.6 cd/m^2^. For measurements in darkness, the displays were switched off. Visual stimuli consisted of sine wave gratings, homogeneous grayscale images and dotted patterns (dot radius 1°, center-to-center distance 5.35°). All stimulus patterns were computer generated and corrected for perspective distortion by projection onto a virtual sphere centered on the animal’s head.

#### Eye movement recordings

Eye movements were measured using an infra-red video tracking system (JAI RM-6740CL monochrome CCD camera, 200 fps). In a few instances (Fig 1A – C) pilocarpine (2%) eye drops were used to reduce pupil dilatation. 2D eye position was computed from the relative distance between pupil center and corneal reflections of the infra-red LEDs (29) and pupil size (30, 31). Epochs containing saccades, eye blinks and motion artifacts were excluded from analysis. Eye velocity was smoothed using a Gaussian smoothing kernel with a SD of 7.5 ms (25 Hz cutoff). For monocular visual stimulation (Fig 1D, 3Aiv, 3Biv, 3Ciii), a miniature blackout cap was placed over the contralateral eye.

#### Intraocular injections

Intravitreal injections were administered under brief isoflurane/O_2_ sedation and topical anesthesia applied to the injection spot (0.4% oxybuprocaine). The eye movements were measured after the mouse recovered completely from the anesthesia. We used injection volumes ranging between 1-3 μL. To block AMPA receptors we used a either 6, 7-dinitroquinoxaline-2,3-dione (DNQX, Tocris, 50 μM final concentration) or cyano-7-nitroquinoxaline-2, 3-dione (CNQX, Tocris, 50 μM final concentration). CNQX and DQNX were assumed to act equally. To block NMDA receptors we used D- (-)-2-Amino-5-phosphonopentanoic acid (D-AP5, Tocris, 10 μM final concentration). To block glycinergic inhibition, we used strychnine (STR, 10 μM final concentration). To block the M-type K-current, we used linopiridine hydrochloride (LP, Tocris, 30 μM inal concentration) Drugs were dissolved in Hank’s Balanced Salt Solution (Sigma-Aldrich).

#### Analysis of eye movement data

Power spectral densities (PSDs) were computed from angular eye velocity using Welch’s method with 4 s window length, 75% overlap between windows and a Hann window function. The fundamental frequency was computed by weighted averaging the frequencies whose magnitude were > 90% of the maximum power in the PSD. For comparisons between different stimulus conditions in *nob* mice, PSDs were normalized to the average power ≤ 2 Hz.

### *In vivo* optic nerve recordings

#### Surgical preparation

All surgical procedures were performed at light adapted levels and have been published previously (32, 33). Briefly, anesthesia was induced with an intraperitoneal injection of a Ringer’s solution containing ketamine and xylazine (127 and 12 mg/kg). Anesthesia was maintained with supplemental subcutaneous injections (~ every 45 min) of anesthetic at 50% of initial concentration. The head was secured in a stereotaxic frame (David Kopf Instruments, Tujunga, CA), a craniotomy was performed and the overlying cortex was removed to expose the optic nerve. Throughout the experiment, body temperature was maintained at 37°C with a feedback controlled heating pad (TC-1000; CWE, Inc., Ardmore, PA). Topical phenylephrine hydrochloride (2.5%) and Tropicamide (1%) ophthalmic solutions were applied to dilate the pupils and paralyze accommodation. Clear zero-powered lenses (34) moistened with artificial tears kept the cornea from drying. At the end of the experiment, animals were euthanized with an overdose of anesthetic followed by cervical dislocation.

#### Extracellular GC axon recordings

Action potentials were recorded extracellularly from single optic nerve axons using sharpened tungsten microelectrodes (impedance = 30-100 MΩ). A reference electrode was inserted subcutaneously. Action potentials from single GC axons were isolated, amplified (X3+Cell; FHC, Bowdoinham, ME), digitized at 15 kHz (Power1401, CED, UK) and stored for offline analysis. Isolated spike trains were simultaneously displayed on an oscilloscope and computer monitor and played over an audio monitor to obtain direct feedback of the cell’s response to visual stimuli. Responses were analyzed offline using Spike2 software v4.24 (Cambridge Electronic Design, USA). Spikes were accumulated within a 50 ms bin width and displayed as post-stimulus time histograms (PSTHs). Each average PSTH was smoothed by fitting it with a raised cosine function with a 50 ms smoothing interval to minimize alteration of the peak firing rate and maximize signal-to-noise ratio (32).

#### Recording and analysis of GC spontaneous activity

Spontaneous activity was determined for GCs in the dark over durations up to 200 s. The presence of a rhythmic component in the spontaneous activity was assessed using a Fast Fourier Transform (FFT) that produced a power spectrum (Spike2 4.24; Cambridge Electronic Design, Cambridge, UK). A peak fundamental frequency was considered significant if its power was three standard errors (SE) above the mean power between 0.5 – 30 Hz (35). To estimate the consistency of the rhythmic activity, the recorded spontaneous activity was divided into segments of 20 s, and FFTs performed on each segment to identify the fundamental peak. The mean fundamental frequency and its standard error were computed across all segments.

### Retinal In vitro GC recordings

Mice were dark-adapted for at least one hour, euthanized with a mixture of CO_2_/O_2_ and cervically dislocated. Under dim red light the eyes were removed and placed in oxygenated Ames medium. The eyecup was prepared by removing the cornea, lens, and as much vitreous humor as possible. Using a fine forceps, the retina was carefully dissected away from the sclera. Small incisions were made to flat-mount the retina. Regardless of whether the isolated retinas were used in the MEA recordings or for whole cell patch clamp recordings, they were continuously superfused with Ames medium or Ringer’s solution and gassed with a mixture of O_2_ and CO_2_ at pH of 7.4 and 29-36°C.

#### Solutions

Ames’ media supplemented with 1.9 g/L of NaHCO3was used as the external bath solution and whole cell patch pipette solution contained [mM]: 112 Cs-Methanesulfonate, 8 CsCl, 10 EGTA, 10 HEPES, 2 ATP-Mg, 0.3 GTP-Na3, pH adjusted to 7.2 with CsOH, ECl =-69.92 mV, except for data shown in Fig 3iii. Here we used a bicarbonate buffered Ringer’s bath solution [mM]: 125 NaCl, 2.5 KCl, 1 MgCl2, 1.25 NaH2PO4, 20 glucose, 26 NaHCO3 and 2 CaCl2 and a whole cell patch pipette solution containing [mM] 120 Cs-gluconate, 1 CaCl2, 1 MgCl2, 10 Na-HEPES, 11 EGTA, 4 ATP & 1 GTP, and 1% LY. In these experiments freshly prepared 6-Cyano-7-nitroquinoxaline-2, 3-dione (CNQX, Tocris) or DNQX (Tocris), D- (-)-2-Amino-5-phosphonopentanoic acid (D-AP5, Tocris) and STR were added to the bath solution to block AMPA, NMDA and glycine receptors, respectively. CNQX and DQNX were assumed to act equally. A cocktail of 50 μM DNQX and 10 μM D-AP5 in the external bath solution was prepared freshly before the experiment. For the dose-response-curve of STR following concentrations were used: 10, 20, 50 and 100 μM STR. Chemicals were obtained from Sigma (St Louis, MS) unless otherwise indicated.

#### Multi-electrode recordings

Isolated retina were placed photoreceptor side up on a perforated 60 electrode array (60pMEA200/30iR-Ti using a MEA2100 system: Multichannel systems, Reutlingen, Germany) in a recording chamber mounted on an Nikon Optiphot-2 upright microscope and viewed under IR with an Olympus 2x objective and video camera (Abus TVCC 20530). Extracellular multi-unit ganglion cell activity was recorded at 25 kHz in MC rack (Multichannel systems, Reutlingen, Germany), zero-phase band-pass filtered (250-6250 Hz) with a 4^th^ order Butterworth filter in Matlab (MathWorks Inc., Natick, MA) and sorted into single unit activity with ‘offline spike sorter’ (Plexon (Dallas, TX). Spikes were detected using an amplitude threshold > 4σ_n_ where σ_n_ is an estimation of the background noise

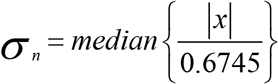

with *x* being the bandpass filtered signal (36). The detected spikes were manually sorted into single units based on principle component or amplitude variables vs time. The clustering vs time approach allowed us to track changes in extracted features of single units occurring over extended recording periods and with differing firing rates

##### Optical stimulator

Light stimuli were generated with Psychophysics Toolbox Version 3 (31). Stimuli were projected onto the retina from the photoreceptor side by a DLP projector (Light Crafter 4500, Wintech, Carsbad, CA) using a custom built 2x water immersion objective. One pixel of the DLP-projector had a size on the retinal surface of 2.1 μm. Only white light stimuli were used. The “dark” light intensity was 6 μW/m^2^ and the maximal “light” intensity was 176.2 μW/m^2^.

##### Analysis of MEA data

For MEA recordings, rhythmic components in the spontaneous activity were assessed using single unit activity during 600 s of darkness. Spike trains were binned into 1 ms intervals and then divided into 5 s non-overlapping periods. Each segment was baseline subtracted, the power spectral density and autocorrelation were evaluated over 120 segments which were then averaged. Autocorrelation were normalized such that they were equal to 1 at zero lag. Power spectral density was estimated by Welch’s modified periodogram using 4 s windows, with 75% overlap, multiplied by a Hamming window function, and a 4 s discrete Fourier transform length.

Synchronized firing between units was assessed by cross-correlation after stimulation a full field light stimulus presented for 500 ms. The flash was preceded and followed by a 500 and 1000 ms period of darkness. Individual unit spike trains were binned into 5 ms interval and the response to 100 repetitions averaged. The mean responses were divided into 250 ms windows, detrended then cross-correlated. For short-time cross-correlations, a 250 ms window advanced in 25 ms steps.

#### Voltage Clamp Measurements of Ganglion Cells

Whole-cell voltage clamp recordings were performed from GFP-labeled ganglion cells in retinas mounted (GC side up) in a recording chamber (Warner Instruments, Hamden, CT, USA). The recording chambers was mounted on a Nikon Eclipse E2000FN microscope (Nikon, Japan) and viewed with a Nikon 60X water immersion objective with infrared differential interference contrast and a video camera (Philips, The Netherlands). The GFP-labeled cells were identified using a short flash of UV light.

##### Optical stimulator

Light stimuli were generated with Psychophysics Toolbox Version 3 (31). Light stimuli were projected onto the retina from the photoreceptor side by an Acer C20 picoprojector via the microscope condenser. One pixel of the projector had a width of 4.8 μm on the retinal surface. The “dark” light intensity was 0.3 cd/m^2^ and the maximal “light” intensity was 450 cd/m^2^.

##### Recording equipment

Whole cell data was recorded with a HEKA EPC10 patch clamp amplifier using PatchMaster software. The data was sampled at 10 kHz and filtered at 5 kHz with a four pole Bessel low pass filter. In a second setup data were collected with a Multiclamp 700B amplifier using Digidata 1322A digitizer (MDS Analytical Technologies, Union City, CA) and Clampex 10.2 software (MDS Analytical Technologies, Union City, CA) to generate command outputs, trigger the light stimulus and acquire and analyze analog whole cell voltages. These data were sampled at 10 kHz and filtered at 2.4 kHz with a four-pole Bessel low pass filter. Matlab (MathWorks Inc., Natick, MA) and Igor.pro (WaveMetrics, Portland, OR, USA) were used to analyze the data.

#### Statistics

Statistical analyses were performed using OriginPro 8 (Northampton, MA, USA) or Matlab (MathWorks Inc., Natick, MA). All mean values are presented ± SEM. Statistical significance was tested using a (paired) Students t-test, or a one way ANOVA. Normality was tested using Shapiro-Wilk test. The distribution of the frequencies of the eye movements (df = 17, p = 0.28), the GCs optic nerve recordings (df = 46, p = 0.71) and the GC excitatory currents (df = 38, p = 0.26) were normally distributed. Differences with p < 0.05 were considered statistically significant.

## Data Availability

The data that support the findings of this study are available from the corresponding author upon reasonable request.

## Acknowledgements

We would like to thank Wim de Graaff for his technical support of this project.

## Supplemental data

**Figure S1.**
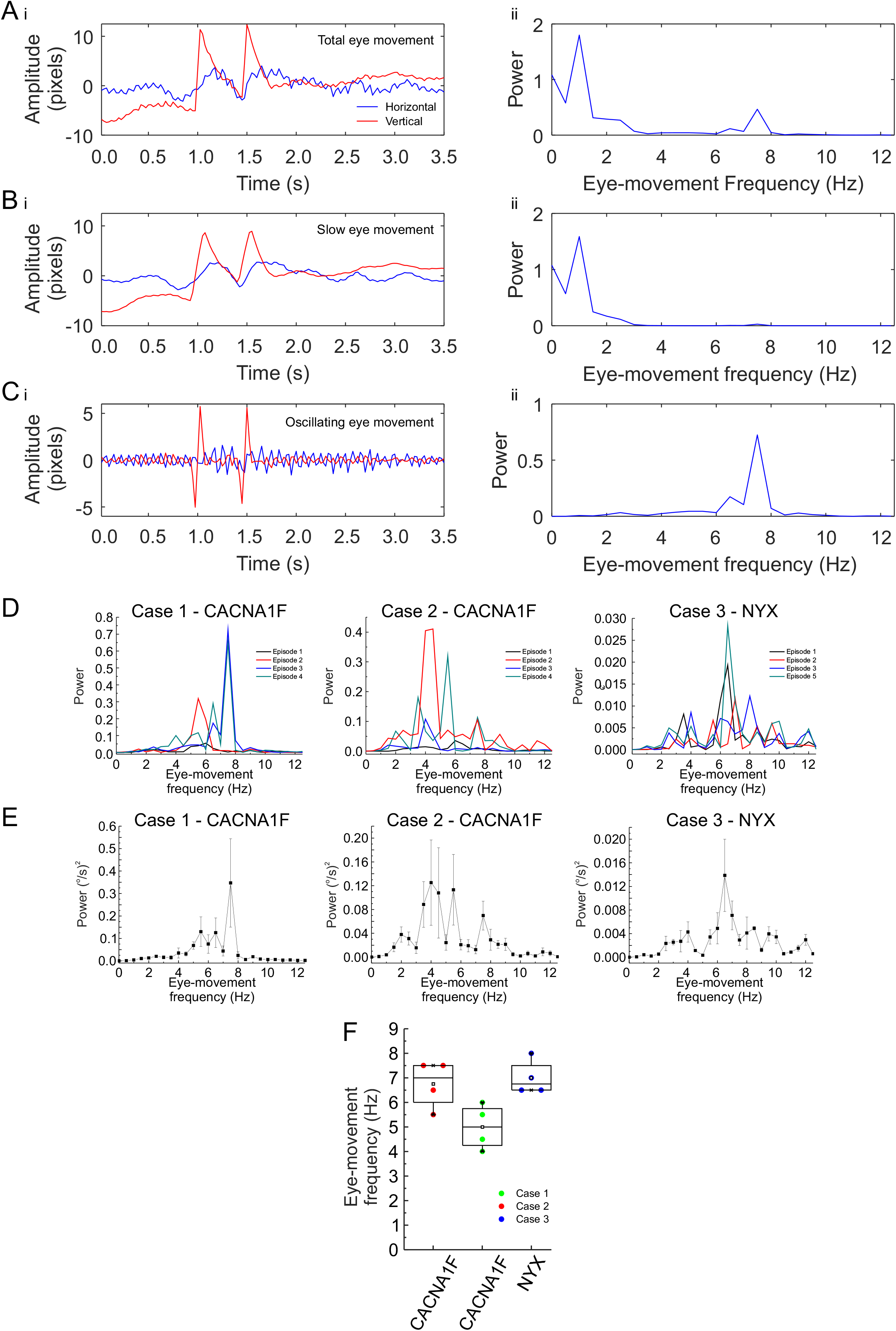
Eyes of CSNB patients oscillate horizontally at 6 Hz. The patients studied by Simonsz et al. (3) had pendular nystagmus combined with tonic downgaze. Although the tonic down gaze disappeared at 2-3 years of age, the horizontal nystagmus remained. We quantified the nystagmus of three patients whose video material was of sufficient quality. The original VHS video movies were digitized, the digital clips were stabilized and aligned using the compositing program Nuke (The Foundry, UK). The nasal corner of the eye and the surrounding area were used as references for the stabilization. After stabilization of the movie, a circle with a fixed diameter the size of the iris was fitted through the iris (See supplementary video). The coordinates of the center of the circle were used as measure for the eye position and the resulting time series of eye positions were Fourier transformed using Matlab. No attempt was made to calibrate the amplitude of the oscillations and the eye movement amplitudes are given in pixels. **Ai)** Total horizontal (blue) and vertical (red) eye movements of a patient #1 in pixels. Large amplitude horizontal and vertical eye movements were present because the children were not fixating and could shift their gaze voluntary. On top of these large amplitude eye movements small amplitude oscillations are visible only for the horizontal eye movements, which is also evident in the power spectrum (**Aii**). There are two peaks visible in the power spectrum: one broad peak below 2 Hz and one narrow peak around 7 Hz. The low frequency component results from gaze shifting. These two components were separated by calculating the moving average over a window of 5 frames (**Bi**) and subtracting this average from the original traces (**Ci**). The power spectra of these two components is shown in **Bii** and **Cii**. For each patient 4 separate episodes were analyzed and the episodes selected not contain eye blinks. **D)** The power spectra of the horizontal eye movements of the individual episodes per patient. All the traces show peaks in the range of 5-8 Hz. **E)** The mean power spectra per patient. **F)** Box plots of the oscillation frequency for the various video clips for each patient. The oscillation frequencies across the various patients did not differ significantly (ANOVA: F = 3.75, df = 11, p = 0.06) and the mean peak oscillation frequency over all patients was 6.25 ± 0.63 Hz (n=3) (Age range: 3 month to 3 years). This oscillation frequency is considerably lower than the 10-20 Hz reported by Pieh et al. (37) in a patient population that included the same patients as described by Simonsz et al (3, 38) and analyzed in this paper. Since Pieh et al. (37) did not disclose the methods how the eye movements were analyzed, it remains puzzling why their estimated of the eye movement oscillation frequency differs so much from ours. We complied with relevant ethical regulations applicable to clinical eye movement recording at the time. The three video recordings of the children aged 0-3 had been made in 1990 and 1992 in the Kantonsspital St. Gallen, Switzerland as a clinical recording of eye movements in the course of their diagnostic work-up and treatment by their treating ophthalmologist (HJS). Informed consent for publication of the video recordings had previously been obtained from the parents. Renewed permission was obtained from the patients themselves who are now adults.

**Figure S2.**
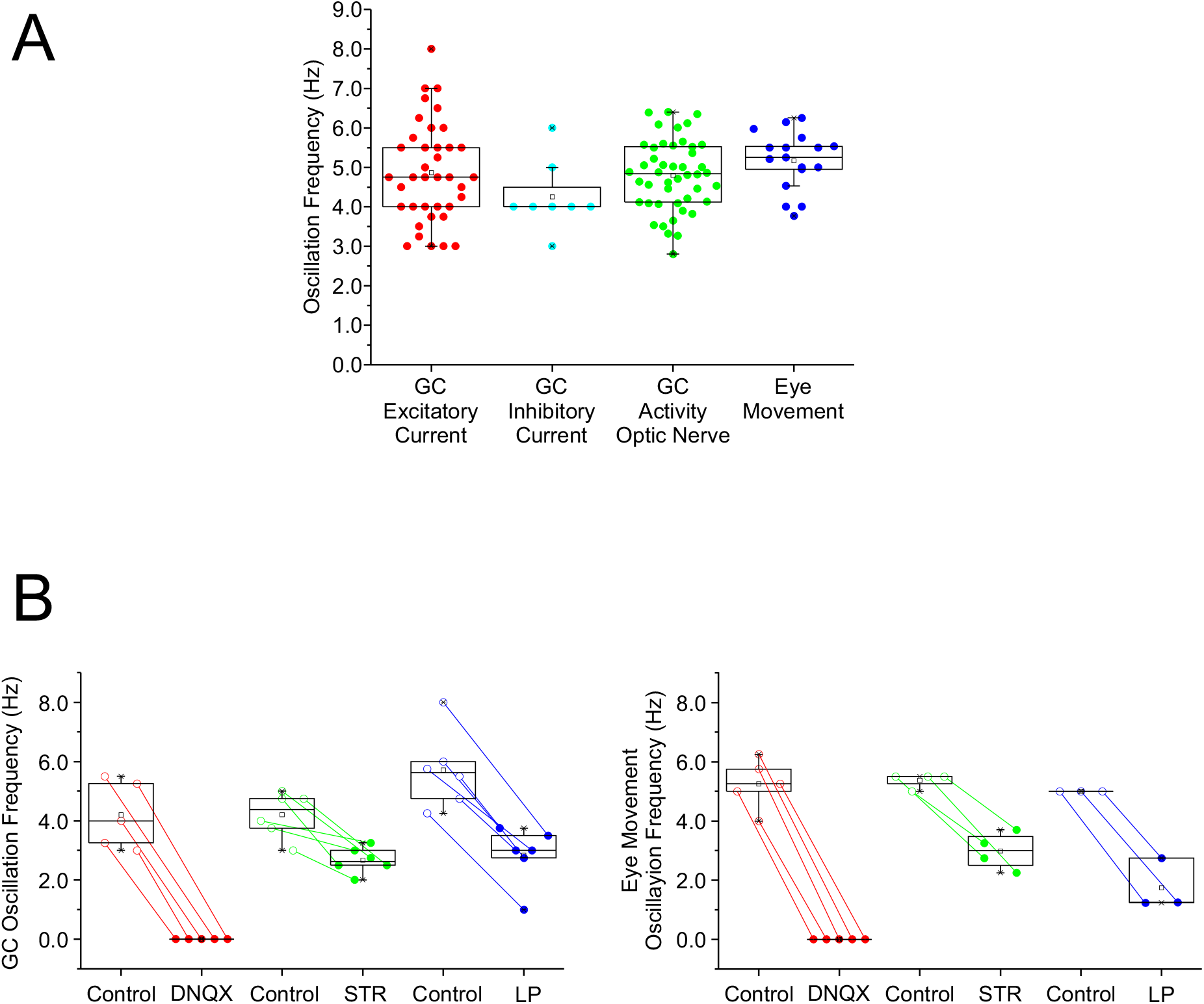
Statistical data of the GC and eye movement oscillation frequencies. The black open squares indicates the mean of all the results, the box indicates the interquartile range (25%-75%), the vertical lines indicate the 5% – 95% range and the colored circles indicate the individual data points. **A)** Box plots of all measurements in control conditions. B) Box plots of all the pharmacological experiments for both the GC oscillations and the eye movements.

**Figure S3.**
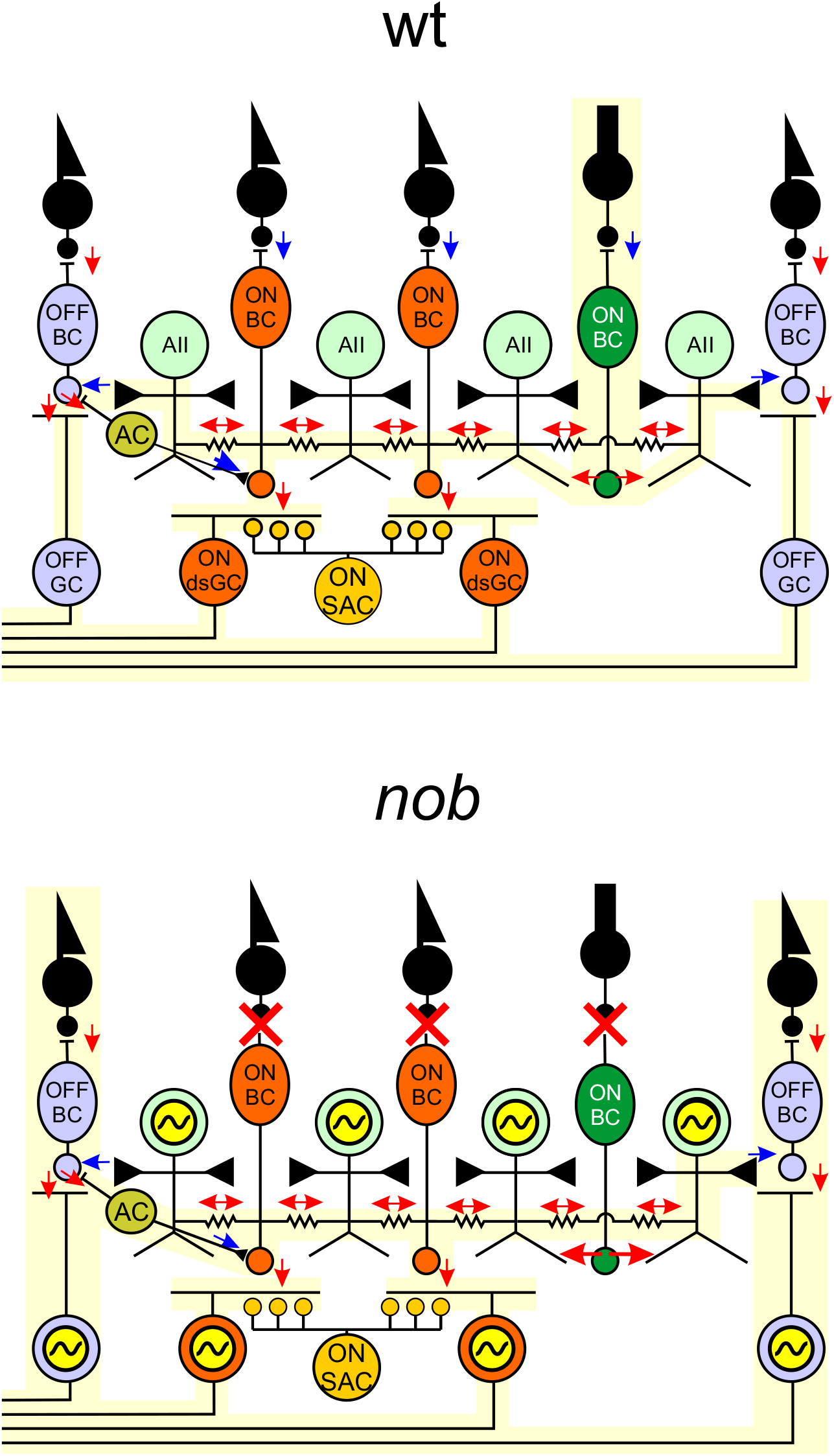
Schematic diagram of the wiring of BCs, AII ACs, starburst-ACs and ON-DSGCs. Wt retinal circuit (top panel) compared to the *nob* retinal circuit (bottom panel). Wt rod ON-BCs provide an excitatory input AII ACs. AII ACs are electrically coupled with each other and to cone ON-BCs and they provide a glycinergic inhibitory input to cone OFF-BCs. Cone ON- and OFF-BCs provide a direct excitatory input to ON- and OFF-GCs. The pathway from rod > rod ON-BC > AII ACs > GCs is the primary rod pathway. AII ACs also receive a glycinergic inhibitory input. ON-DSGCs receive excitatory input from cone ON-BCs and are inhibited by ON-starburst amacrine cells (SACs). The interaction between ON-BC excitation and ON-SACs inhibition produces direction selective responses in ON-DSGCs. OFF-BC excitatory input to AII ACs is relayed to cone ON-BCs via electrical coupling as well as via crossover inhibition between the ON- and OFF-pathways. In *nob* retina, signaling from the rod and cone photoreceptors to ON-BCs is lost because the synapse between photoreceptors and ON-BCs is non-functional. The consequence is that ON light responses are absent and that the GCs have oscillating burst spike activity. These oscillations most likely originate in AII ACs just as in *rd1*-mice (39). Interaction between fast sodium and potassium channels in AII ACs, in combination with a slow potassium channel can lead to a bursting spiking behavior with a burst frequency dependent on the AII AC membrane potential (39). In *rd1*-mice, the input to AII ACs is lost with rod and cone degeneration, leading to a change in AII AC membrane potential, which induces oscillations at about 10 Hz. In *nob* mice, AII ACs lose only their ON-input. We hypothesize that as a result the AII AC membrane potential will not shift as far as in rd1 mice and the AII ACs oscillations will be slower (5 Hz). The *nob* AII ACs transmit this oscillating signal back to the BC-synaptic terminals and to the GCs, which induces 5 Hz oscillations in both the excitatory and inhibitory inputs of the GCs. Since the sign of the AII AC inputs is opposite in ON-BCs and OFF-BCs, the *nob* ON- and OFF-GCs oscillate in antiphase. In the dark the AII ACs might oscillate independent of each other. A global light stimulus will reset all AII ACs and, in that way, synchronize them.

